# Rho-associated kinase regulates Langerhans cell morphology and responsiveness to tissue damage

**DOI:** 10.1101/2023.07.28.550974

**Authors:** Eric Peterman, Elgene J.A. Quitevis, Camille E.A. Goo, Jeffrey P. Rasmussen

**Author notes:** Correspondence (E.P.); (J.P.R.).

## Abstract

Skin is often the first physical barrier to encounter invading pathogens and physical damage. Damage to the skin must be resolved quickly and efficiently to maintain organ homeostasis. Epidermal-resident immune cells known as Langerhans cells use dendritic protrusions to dynamically surveil the skin microenvironment, which contains epithelial keratinocytes and somatosensory peripheral axons. The mechanisms governing Langerhans cell dendrite dynamics and responses to tissue damage are not well understood. Using skin explants from adult zebrafish, we show that Langerhans cells maintain normal surveillance activity following axonal degeneration and use their dynamic dendrites to engulf small axonal debris. By contrast, a ramified-to-rounded shape transition accommodates the engulfment of larger keratinocyte debris. We find that Langerhans cell dendrites are richly populated with actin and sensitive to a broad spectrum actin inhibitor. We further show that Rho-associated kinase (ROCK) inhibition leads to elongated dendrites, perturbed clearance of large debris, and reduced Langerhans cell migration to tissue-scale wounds. Altogether, our work describes the unique dynamics of Langerhans cells and involvement of the ROCK pathway in immune cell responses to damage of varying magnitude.

## INTRODUCTION

Squamous epithelia coat external interfaces, including skin and mucocutaneous organs, and provide essential environmental barriers. Due to their close proximity to the environment, these epithelia are frequently damaged. To restore barrier function after damage, surrounding epithelial and immune cells mount coordinated efforts to close the wound and eliminate cellular debris to prevent chronic inflammation, respectively.^1–3^ The confined nature of epithelial tissues presents challenges for resident-immune cells to infiltrate the wound and clear debris.

Skin harbors dense networks of epithelial keratinocytes, somatosensory peripheral axons, and immune cell types. The outermost layer of skin, the epidermis, contains immune cells known as Langerhans cells essential to the wound healing response.^4^ Intriguingly, Langerhans cells display a unique mixture of dendritic cell and macrophage properties.^5^ Historically, Langerhans cells have been mainly studied for their dendritic cell capabilities: intercepting pathogens and antigens to promote adaptive immune responses following emigration from the skin to lymph nodes.^5–7^ However, recent work shows that Langerhans cells share origins and a genetic dependence on IL-34/Csf1r signaling with tissue-resident macrophages in other organs.^8–11^ Additionally, live-cell imaging studies in zebrafish have identified macrophage-like roles for Langerhans cells locally within the epidermis, including engulfment of degenerating axonal debris and migration to sites of keratinocyte damage.^12, 13^ The mechanisms of how Langerhans cells quickly and efficiently respond to tissue damage are unknown.

Langerhans cells extend thin membrane protrusions known as dendrites between neighboring keratinocytes and in close proximity to somatosensory axons.^14–20^ This dendritic morphology allows Langerhans cells to surveil large areas of the epidermis. Langerhans cell dendrites are dynamic structures^15, 21^ proposed to regulate the regular positioning of Langerhans cells within the epidermis and their uptake of external antigens and pathogens.^19, 20^ What controls Langerhans cell dendrite dynamics and morphogenesis? Loss of E-cadherin, a major linkage between the plasma membrane and actin cytoskeleton, results in decreased Langerhans cell dendrites.^22^ Consistent with a role for the actin cytoskeleton in dendrite morphogenesis, genetic deletion of the Rho family GTPases Cdc42 or Rac1, which regulate actin remodeling, reduces Langerhans cell dendritic branching.^20, 23^ Despite these advances, the cytoskeletal control of Langerhans cell dendrite dynamics remains poorly understood.

The optical accessibility of zebrafish skin provides an attractive experimental system to study Langerhans cell dynamics. Similar to mammals, the adult zebrafish epidermis contains dendritic Langerhans cells that intermingle with peripheral somatosensory axons and stratified layers of keratinocytes **(Figure 1A)**.^11–13, 24–26^ In this study, we use the genetic and imaging advantages of zebrafish along with our previously established skin explant assay to better understand Langerhans cell dendrite morphogenesis, dynamics, and responses to several forms of tissue damage.

**Figure 1.**
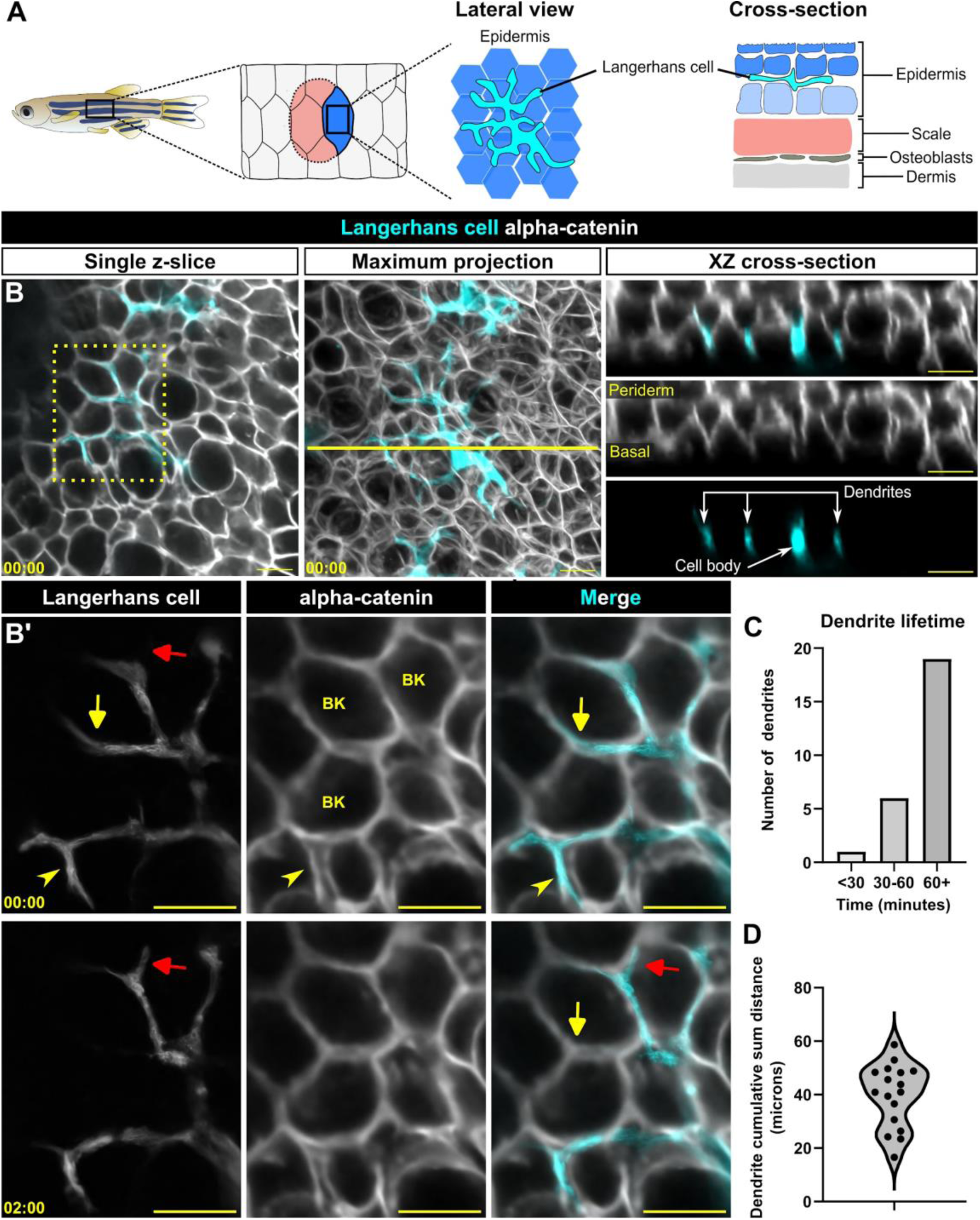
Langerhans cell dendrites survey the epidermis in the absence of stimulus. **A.** Schematic of adult zebrafish trunk skin, showing lateral and cross-section views of the epidermis. **B.** Single z-slice (left), maximum intensity projection (middle) and YZ projection along yellow line in middle panel (right) images of *Tg(mpeg1:NTR-EYFP)*-positive cells and *Gt(ctnna1-Citrine)* labeled epidermal cell membranes, illustrating the complexity of skin epidermis. **B’.** Inset from dashed box in B (single z-slice), illustrating Langerhans cell dendrite retraction. See also Supplemental Video 1. Yellow arrow denotes dendrite retraction over time, red arrow denotes dendrite extension over time, arrowhead denotes a dendrite occupying space between keratinocytes. BK, basal keratinocyte. **C.** Quantification of dendrite lifetime, n = 26 dendrites tracked from 7 cells. **D.** Violin plot of cumulative sum distance traveled by dendrites over a 10 minute window, n = 17 dendrites tracked from 7 cells. Timestamps in **(B, B’)** denote mm:ss. Scale bars in **(B, B’)** denote 10 microns.

## RESULTS

### Langerhans cell dendrites are long-lived yet highly dynamic during homeostasis

At steady-state, the dendrites of murine Langerhans cells cyclically extend and retract.^15, 20, 21^ Previously, we showed that zebrafish Langerhans cells engulf axonal debris following cutaneous axon degeneration via their dynamic dendrites.^13^ To understand the basis for these dendrite dynamics in more detail, we used our established skin explant assay^13^ to monitor Langerhans cell dendrite motility by collecting confocal z-stacks every 30 seconds. Zebrafish Langerhans cells express transgenes driven by the *mpeg1.1* promoter.^11, 12, 25^ We co-imaged Langerhans cells expressing a cytoplasmic reporter (*Tg(mpeg1:NTR-EYFP)*)^27^ and epidermal cell junctions labeled by an *alpha-catenin* (*ctnna1*) gene trap reporter.^28^ We found that Langerhans cell dendrites extended and retracted between keratinocyte membranes (**Figure 1B,B’, Supplemental Video 1**). We skeletonized Langerhans cells **(Supplemental Figure 1A)** and measured dendrite lifetimes and displacements. Most primary dendrites had relatively long lifetimes (> 60 min) **(Figure 1C).** Remarkably, some dendrites extended and retracted cumulative distances of up to 60 microns within a 10 minute frame **(Figure 1D)**. Overall, our observations indicate that zebrafish Langerhans cell dendrites are long-lived and motile in the absence of stimulus, consistent with previous work in mice.^21^

### Langerhans cell dendrite morphology does not change following axon degeneration

Following skin explant, somatosensory axons in the skin undergo a stereotypical degeneration process known as Wallerian degeneration, resulting in large quantities of debris internalized by Langerhans cells.^13^ We questioned if axon degeneration altered Langerhans cell dendrite number, surface area, or cell morphology. To track somatosensory axon debris following Wallerian degeneration, we used *Tg(p2rx3a:lexA;LexAOP:mCherry)*^29^ (hereafter referred to as *Tg(p2rx3a:mCherry)*), a reporter expressed in skin-innervating adult dorsal root ganglion neurons.^30^ Interestingly, we found that Langerhans cell dendrite number was unchanged within the first 60 minutes after axon degeneration, a period in which Langerhans cells actively internalize axonal debris^13^ **(Figure 2A–C, Supplemental Video 2)**. To assess Langerhans cell morphology, we calculated cell circularity before and after axon degeneration and found no significant changes **(Figure 2D)**. To assess the surface area covered, we measured the convex hull of Langerhans cells and similarly did not observe a significant change before and after axon degeneration **(Figure 2E)**. Based on these observations, we concluded that axon degeneration did not affect Langerhans cell behavior or morphology within our observation window. This suggests Langerhans cells internalize axonal debris as part of their homeostatic surveillance dynamics.

**Figure 2.**
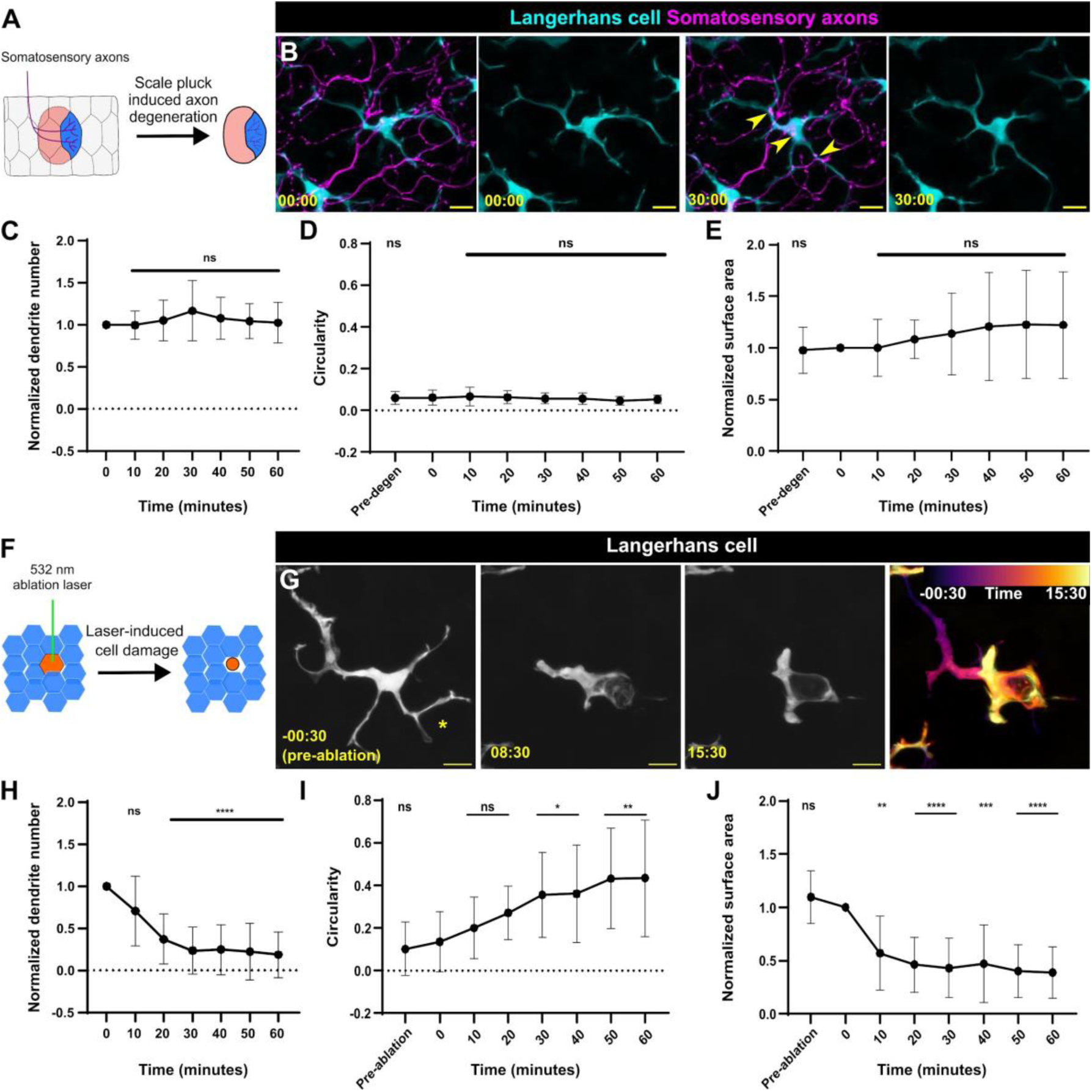
Langerhans cells use injury-dependent engulfment modes. **A.** Schematic illustrating scale removal and subsequent axon degeneration. **B.** Representative images showing *Tg(mpeg1:NTR-EYFP)*-positive Langerhans cell morphology during engulfment of *Tg(p2rx3a:mCherry)*-positive axonal debris. Arrowheads denote internalized axonal debris. **C.** Quantification of dendrite number following axon degeneration. Dendrite number is normalized to the number present at time of axon degeneration. n = 17 cells from N = 12 scales. **D.** Quantification of circularity before and after axon degeneration. n = 16 cells from N = 12 scales. **E.** Quantification of surface area covered (convex hull) before and after axon degeneration. Surface area is normalized to area at time of axon degeneration. n = 15 cells from N = 12 scales. **F.** Schematic illustrating laser-mediated cell damage. **G.** Representative images showing *Tg(mpeg1:NTR-EYFP)-*positive Langerhans cell morphology during engulfment of laser-induced cellular debris. Asterisk denotes site of laser ablation. Far-right panel shows temporal color coding as indicated in the legend depicting shape change as engulfment proceeds. **H.** Quantification of dendrite number following laser ablation. Dendrite number is normalized to the number present at time of laser ablation. n = 10 cells from N = 5 scales. **I.** Quantification of circularity before and after laser ablation. n = 12 cells from N = 5 scales. **J.** Quantification of surface area covered (convex hull) before and after laser ablation. Surface area is normalized to area at time of laser ablation. n = 11 cells from N = 5 scales. * = p < 0.05, ** = p < 0.01, **** = p < 0.0001. One-way ANOVA followed by Bonferroni post-tests were used to determine significance compared to time = 0 at each time point. In **(C-E)** and **(H-J)**, data points represent averages, error bars represent standard deviation. Timestamps in **(B, G)** denotes mm:ss. Scale bars in **(B, G)** denote 10 microns.

### Langerhans cells undergo a ramified-to-rounded shape transition to engulf large cellular debris

While Langerhans cells have been classified as tissue-resident macrophages based on ontogeny,^5^ their phagocytic capabilities are poorly described.^31^ Since phagocytosis depends on properties of the target substrate, including size,^32–34^ and axonal debris is relatively small (∼1 to 3 microns in diameter), we questioned whether Langerhans cells could engulf larger types of cellular debris. To this end, we employed an ablation laser to create reproducible keratinocyte damage, allowing us to assess if Langerhans cells react to and engulf larger debris (∼5 to 20 microns in diameter). In contrast to the apparent lack of reaction to axonal debris, Langerhans cells underwent a rapid, stereotyped series of shape changes to engulf keratinocyte debris that we refer to as a ramified-to-rounded shape transition **(Figure 2F, G, Supplemental Figure 2A, Supplemental Video 3)**. During this process, Langerhans cells retracted dendrites distal from the site of engulfment, leading to fewer dendrites, while one or two proximal dendrites extended toward the cellular debris to facilitate engulfment **(Figure 2H).** As engulfment proceeded, Langerhans cells completed the ramified-to-rounded shape transition by fully surrounding the debris, leading to an increase in circularity and decrease in surface area **(Figure 2I, J)**. We occasionally observed Langerhans cells undergoing similar shape transitions to engulf large pieces of cellular debris in the absence of laser damage, suggesting this shape transition was not due to the laser itself **(Supplemental Figure 2B).** These data establish a reproducible method for monitoring Langerhans cell reactions to local keratinocyte damage. Overall, our results indicate that Langerhans cells proceed through a ramified-to-rounded shape transition to engulf larger pieces of cellular debris, while engulfment of smaller axonal debris requires no shape transition.

### Langerhans cell dendrite motility and debris engulfment require actin

Actin polymerization is required for extending smaller actin-based membrane protrusions such as filopodia and microvilli.^35^ Prior work identified roles for the actin regulators Rac1 and Cdc42 in promoting Langerhans cell dendrite morphogenesis,^20, 23^ suggesting that actin dynamics may regulate dendrite behaviors. However, these analyses were performed days to weeks after genetic deletion, confounding the interpretation. To our knowledge, neither visualization of actin nor acute perturbations of the cytoskeleton in Langerhans cells have been reported. To better understand actin dynamics in Langerhans cells, we created a stable transgenic line *Tg(mpeg1:Lifeact-GFP)*, in which the Lifeact-GFP probe^36^ labels filamentous actin (F-actin) in Langerhans cells. We explanted skin from these fish and imaged every 5 seconds to visualize dendrite dynamics. During dendrite retraction, Lifeact-GFP strongly localized to the distal end of the dendrite **(Figure 3A, Supplemental Video 4)**, suggesting F-actin dynamically reorganizes during dendrite retraction. To examine the necessity for actin dynamics during dendrite motility, we explanted skin from *Tg(mpeg1:Lifeact-GFP)* fish and imaged in the presence of Latrunculin B (LatB), an inhibitor of actin dynamics,^37^ or vehicle control **(Figure 3B)**. Following addition of LatB, dendrite dynamics slowed and dendrite length shortened, resulting in cells covering a smaller surface area compared to controls **(Figure 3C).** Washout of LatB rapidly restored dendrite dynamics and morphology, suggesting its effects were reversible (**Supplemental Figure 3A)**.

**Figure 3.**
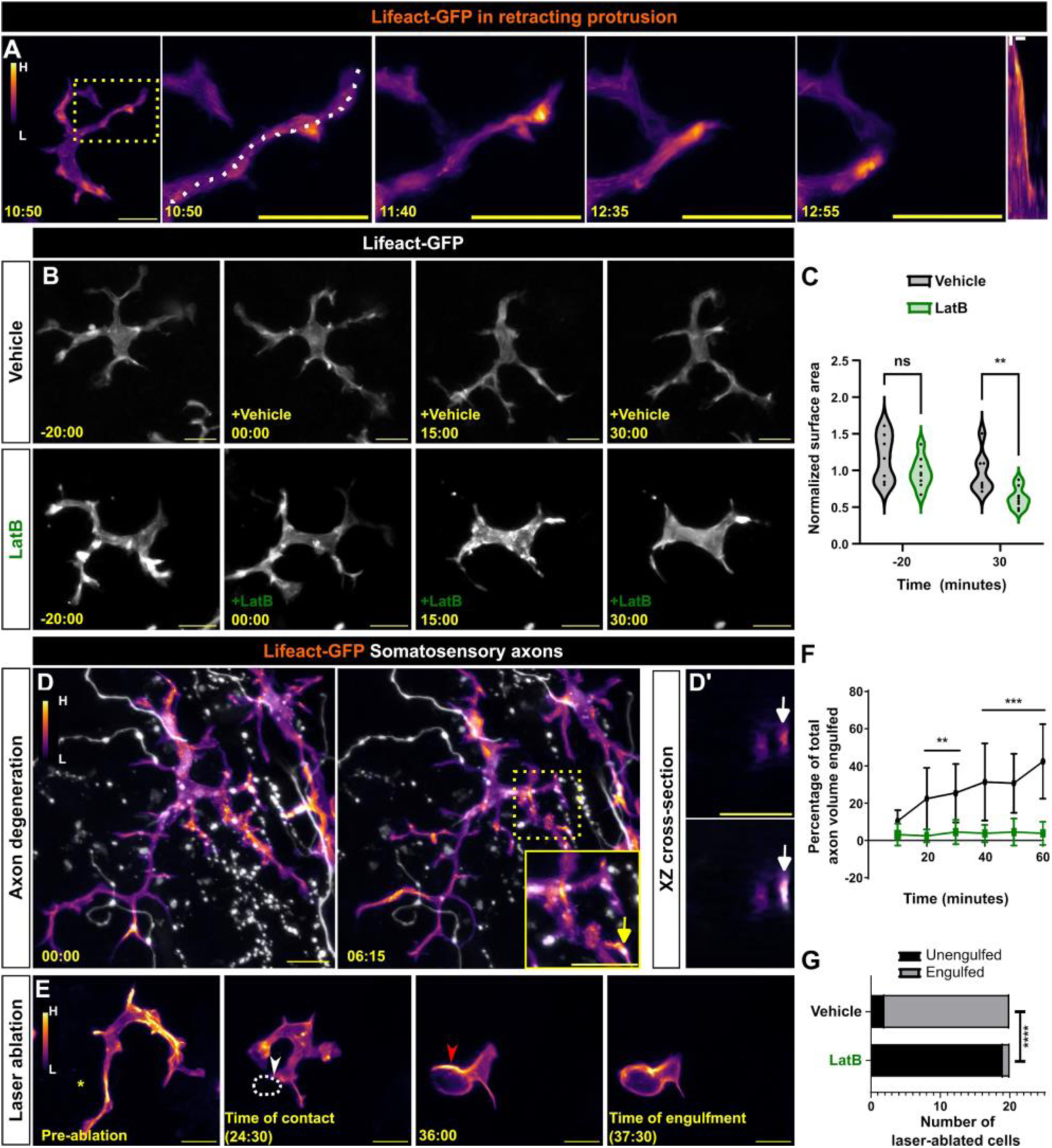
Actin localization and requirements in Langerhans cells during debris engulfment. **A.** Stills of *Tg(mpeg1:Lifeact-GFP)*-positive Langerhans cell depicting Lifeact-GFP localization in a retracting dendrite. Lifeact signal intensity color-coded from low (L) to high (H). Dotted box indicates dendrite of interest that is magnified in panels at right. Dotted line depicts area traced for kymograph (right-most panel). **B.** Stills of *Tg(mpeg1:Lifeact-GFP)*-positive Langerhans cells depicting loss of surface area coverage after Latrunculin B (LatB) treatment, but not after vehicle (ethanol) control. **C.** Violin plots of surface area coverage before and after LatB treatment. n = 7 cells from N = 3 scales for vehicle control, n = 8 cells from N = 3 scales for LatB. **D.** Stills of *Tg(mpeg1:Lifeact-GFP)*-positive Langerhans cell depicting Lifeact-GFP localization at sites of axonal debris engulfment. Dotted box surrounds region of interest and is magnified in inset. Arrows in **(D, inset)** point to the same debris in XZ cross-sections as in **(D’). D’** (top) shows Lifeact-GFP only, **D’** (bottom) shows merge. **E.** Stills of *Tg(mpeg1:Lifeact-GFP)*-positive Langerhans cell depicting Lifeact-GFP localization during engulfment of large cellular debris. Asterisk denotes site of laser ablation, yellow arrowhead denotes site of cell-debris contact, dotted white line denotes cellular debris, red arrowhead denotes actin enrichment during engulfment. **F.** Quantification of axonal debris engulfment in vehicle-treated controls or LatB-treated scales. n = 9 cells from N = 4 scales for control and n = 9 cells from N = 6 scales for LatB. **G.** Quantification of large debris engulfment by vehicle-treated controls or LatB-treated scales, n = 20 ablated cells from N = 2 scales for vehicle and LatB. * = p < 0.05, ** = p < 0.01, *** = p < 0.001, **** = p < 0.0001. Mann-Whitney U test was used to determine significance in **(C)**. Two-way ANOVA followed by Bonferroni post-tests was used to determine significance between groups at each time point in **(F).** Fisher’s exact test was used to determine significance in **(G).** In **(F)**, data points represent averages, error bars represent standard deviation. Timestamps denote mm:ss. Scale bars in **(A, B, D, E)** denote 10 microns, scale bars in **(D, inset, D’)** denote 5 microns, scale bars in **(A, kymograph)** denote 60s (horizontal) and 2 microns (vertical).

To determine F-actin localization during debris engulfment, we triggered axon degeneration by explanting scales from *Tg(mpeg1:Lifeact-GFP);Tg(p2rx3a:mCherry)* double transgenic fish. Upon axon degeneration, we observed Lifeact+ foci within dendrites colocalizing with debris as it was engulfed **(Figure 3D, Supplemental Video 5)**. Similarly, we observed Lifeact-rich phagocytic cup formation during engulfment of larger debris after laser-induced damage **(Figure 3E, Supplemental Video 6)**.

To test if the decrease in dendrite movement and length we observed after LatB treatment correlated to a decrease in debris engulfment, we repeated our engulfment assays in the presence of LatB or vehicle control. After axon degeneration, we recorded a significantly decreased ability for LatB-treated Langerhans cells to engulf axonal debris **(Figure 3F)**. Similarly, using our laser-induced damage paradigm, we found that 18/20 ablated cells were engulfed after 40 minutes in vehicle-treated conditions, whereas only 1/20 ablated cells were engulfed in LatB-treated conditions **(Figure 3G).** Combined, these data suggest that actin dynamics are required for Langerhans cell dendrite maintenance and debris engulfment.

### Rho-associated kinase is required for dendrite morphology and motility

Rho-associated kinase (ROCK) functions downstream of the small GTPase RhoA to regulate actin polymerization. In two-dimensional models of cell migration and chemotaxis, ROCK is required for cells to retract their trailing edge.^38–41^ Therefore, we hypothesized the ROCK pathway may control Langerhans cell dendrite dynamics and/or morphology. To test if ROCK regulated Langerhans cell dendrite dynamics, we treated scale explants with the ROCK inhibitor Y-27632 (referred to as ROCKi).^42^ Following ROCKi treatment, we observed continuous dendrite elongation, which plateaued 60 minutes post-treatment **(Figure 4A, Supplemental Video 7)**. Accompanying this, we recorded an increase in dendrite lifetime and surface area covered **(Figure 4B, C)**. Washout of ROCKi returned cells to normal morphology and cell motility **(Supplemental Figure 3B)**, suggesting its effects were reversible. Furthermore, treating cells with a different ROCK inhibitor (Rockout) recapitulated our results with Y-27632 **(Supplemental Figure 3C)**, indicating dendrite elongation was specific to ROCK inhibition. Imaging of epidermal cell membranes revealed that whole tissue organization was not impacted following ROCKi treatment **(Supplemental Figure 3D)**. We examined another immune cell type present in the skin, *lck+* T cells,^43^ and found that these normally amoeboid cells did not exhibit a change in surface area in the presence of ROCKi **(Supplemental Figure 3E)**. These results suggest ROCK inhibition elongates Langerhans cell dendrites without affecting the morphology of other immune cells or perturbing total tissue integrity.

**Figure 4.**
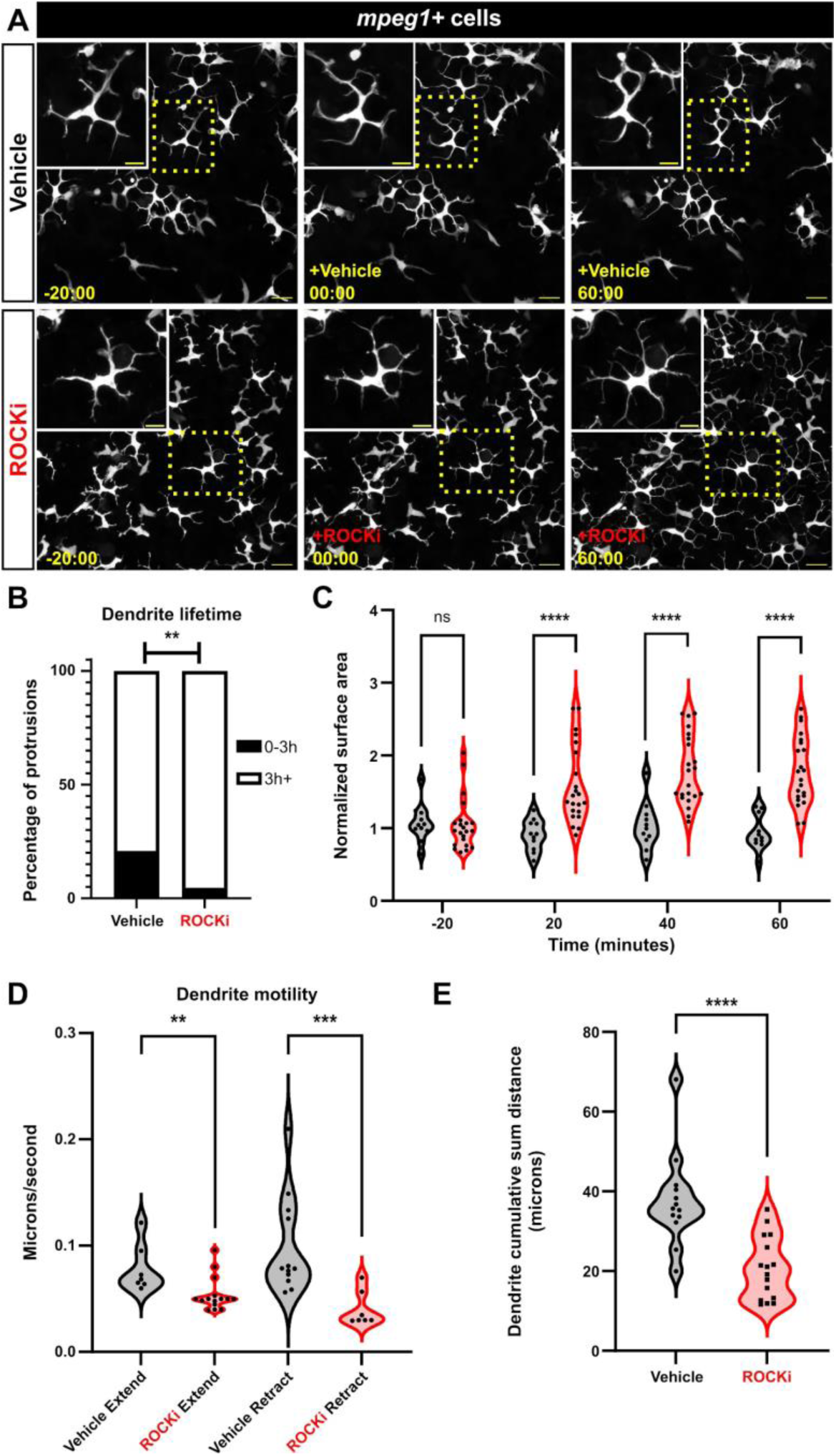
ROCK regulates Langerhans cell dendrite length and dynamics. **A.** Still images showing *Tg(mpeg1:NTR-EYFP)*-positive cells after vehicle treatment (top row) or ROCK inhibition (bottom row). Dotted boxes indicate Langerhans cells magnified in insets. **B.** Quantification of dendrite lifetime after vehicle treatment or ROCK inhibition. n = 91 dendrites from 17 cells, N = 6 scales in vehicle treatment and n = 86 dendrites from 24 cells, N = 5 scales in ROCKi treatment. **C.** Violin plots of surface area covered (convex hull) over time before and after vehicle treatment and ROCK inhibition. n = 11 for vehicle, n = 22 for ROCKi. **D.** Violin plots of dendrite extension and retraction speeds. n = 7 from 4 cells dendrites in vehicle extend, n = 15 from 6 cells in ROCKi extend, n = 12 from 4 cells in vehicle retract, n = 7 from 3 cells in ROCKi retract. **E.** Violin plots of cumulative sum distance traveled over a 10 minute window. n = 13 cells for vehicle control, n = 16 cells for ROCKi. ** = p <0.01, *** = p < 0.001, **** = p < 0.0001. Fisher’s exact test was used to determine significance in **(B)**. Two-way ANOVA followed by Bonferroni post-tests were used to determine significance between groups at each time point in **(C)**. Mann-Whitney U tests were used to determine significance between groups in **(D, E).** Timestamps in **(A)** denotes mm:ss. Scale bars in **(A)** denote 20 microns, scale bars in **(A, inset)** denote 10 microns.

Since ROCK is required for cellular dynamics in other contexts,^44^ we next measured Langerhans cell dendrite motility by quantifying extension and retraction speeds. We found that ROCK inhibition led to a moderate, but significant, decrease in dendrite extension speed. And, while dendrite retraction was relatively rare following ROCKi treatment, we found that retraction speed was significantly decreased **(Figure 4D)**. Consistent with the observation that extension and retraction speeds were altered, we found that dendrites in ROCKi-treated conditions traveled smaller cumulative sum distances over a 10 minute period in comparison to vehicle-treated controls **(Figure 4E)**. Combined, these data suggest that ROCK dictates the homeostatic surveillance of Langerhans cells by controlling dendrite growth and dynamics.

ROCK has numerous substrates, many of which are involved in the actin cytoskeleton’s roles in controlling cell contraction, polarity, and migration.^44^ One downstream effector of ROCK is non-muscle myosin-II (NMII), a regulator of actomyosin contractility. To visualize NMII during protrusion retraction, we used *Tg(actb2:myl12.1-EGFP)*,^45^ which expresses a myosin light chain fused to EGFP under the ubiquitous *actb2* promoter. Interestingly, although we observed localized NMII in retracting dendrites, Langerhans cell surface area did not increase after NMII inhibition with blebbistatin **(Supplemental Figure 4A-C).** Overactivation of NMII via inhibition of myosin phosphatase with Calyculin A^46^ resulted in transient and moderate dendrite retraction **(Supplemental Figure 4D).** These data suggest that activation of NMII likely promotes dendrite retraction, but additional pathways downstream of ROCK may regulate Langerhans cell morphology.

### ROCK promotes Langerhans cell shape transition and migration to wounds

What are the functional consequences of ROCK inhibition on Langerhans cell responses to epidermal damage? ROCK is specifically required for phagocytosis^47–49^ and perturbing ROCK alters macrophage motility and environmental sampling *in vitro*.^50^ To assess a role for ROCK in small debris engulfment, we removed scales from *Tg(mpeg1:NTR-EYFP);Tg(p2rx3a:mCherry)* fish and treated them with ROCKi. After axon degeneration, we recorded little change in the ability of ROCKi-treated Langerhans cells to engulf axonal debris compared to controls **(Figure 5A, B).** By contrast, ROCKi treatment perturbed the ability of Langerhans cells to complete the ramified-to-rounded shape transition following laser-induced cellular damage. Specifically, ROCKi-treated cells did not retract trailing dendrites as readily as controls, leading to significantly decreased circularity and increased surface area at time of engulfment **(Figure 5C-E, Supplemental Video 8)**. Remarkably, despite this altered morphology, ROCKi-treated Langerhans cells still reacted to damage, with moderately increased time until first contact when compared to controls **(Figure 5F)**. Finally, we found that ROCKi treatment significantly increased the time required for phagocytic cup closure **(Figure 5G)**. Thus, we conclude that ROCK promotes the ramified-to-rounded shape transition that accompanies engulfment of large debris.

**Figure 5.**
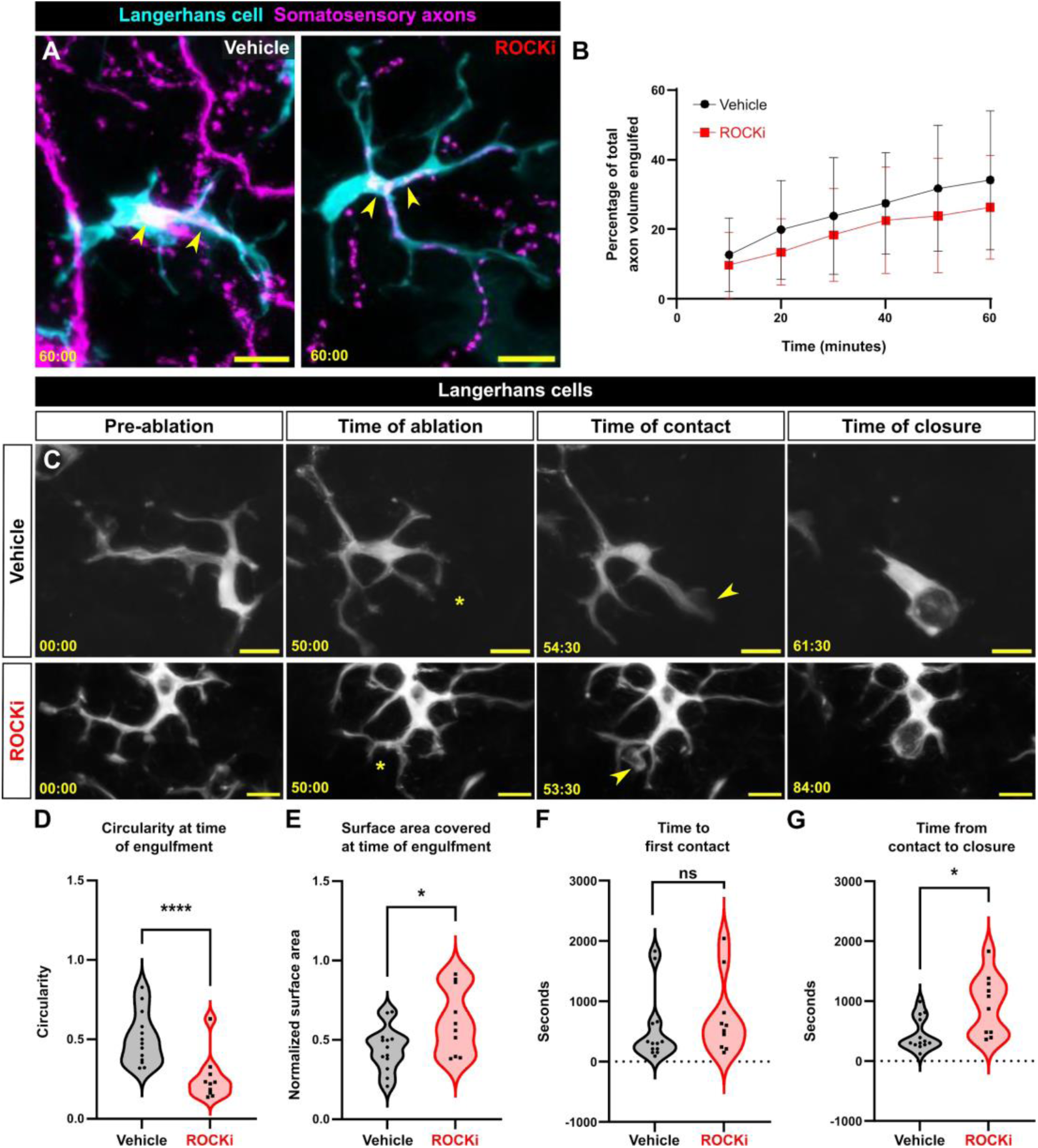
Effects of ROCK inhibition on engulfment of axonal and keratinocyte debris. **A.** Still images showing *Tg(mpeg1:NTR-EYFP)*-positive Langerhans cell engulfing *Tg(p2rx3a:mCherry)*-positive axonal debris after vehicle treatment or ROCK inhibition. Yellow arrowheads indicate engulfed axonal debris. **B.** Quantification of axonal volume engulfed in vehicle or ROCKi conditions. n = 12 cells from N = 10 scales for vehicle, n = 16 cells from N = 9 scales for ROCKi. **C.** Still images showing *Tg(mpeg1:NTR-EYFP)*-positive Langerhans cells after vehicle treatment (top row) or ROCK inhibition (bottom row) in the context of cell ablation. Asterisks denote sites of laser ablation. Arrowheads denote sites of contact with debris. **D.** Violin plots of circularity at time of engulfment. n = 13 cells, N = 3 scales for vehicle and n = 15 cells, N = 4 scales for ROCKi. **E.** Violin plots of surface area covered at time of engulfment. n = 13 cells for vehicle, N = 3 scales and n = 15 cells, N = 4 scales for ROCKi. **F.** Violin plots of the amount of time from ablation to first contact of debris. n = 13 cells, N = 3 scales for vehicle and n = 15 cells, N = 4 scales for ROCKi. **G.** Violin plots of the amount of time from first contact of debris to closure of phagocytic cup. n = 13 cells, N = 3 scales for vehicle and n = 15 cells, N = 4 scales for ROCKi. * = p < 0.05, ** = p <0.01. Two-way ANOVA followed by Bonferroni post-tests was used to determine significance between groups at each time point in **(B).** Mann-Whitney U tests were used to determine significance in **(D-G).** In **(B)**, data points represent averages, error bars represent standard deviation. Timestamps in **(A, C)** denote mm:ss. Scale bars in **(A, C)** denote 10 microns.

Previous work found that zebrafish Langerhans cells migrate to epidermal scratch wounds.^12^ To examine if Langerhans cell responses to tissue-scale wounds required ROCK, we treated explanted scales with vehicle or ROCKi and introduced a large (>10,000 μm^2^) epidermal wound via mechanical injury **(Figure 6A)**. Owing to impaired cell motility, we observed a significant decrease in the number of Langerhans cells in the wound margin following ROCKi treatment **(Figure 6B,C, Supplemental Video 9)**. These results show that Langerhans cells require ROCK for efficient migration to epidermal wounds, and altogether, that ROCK promotes dendrite dynamics and responses to specific types of tissue damage.

**Figure 6.**
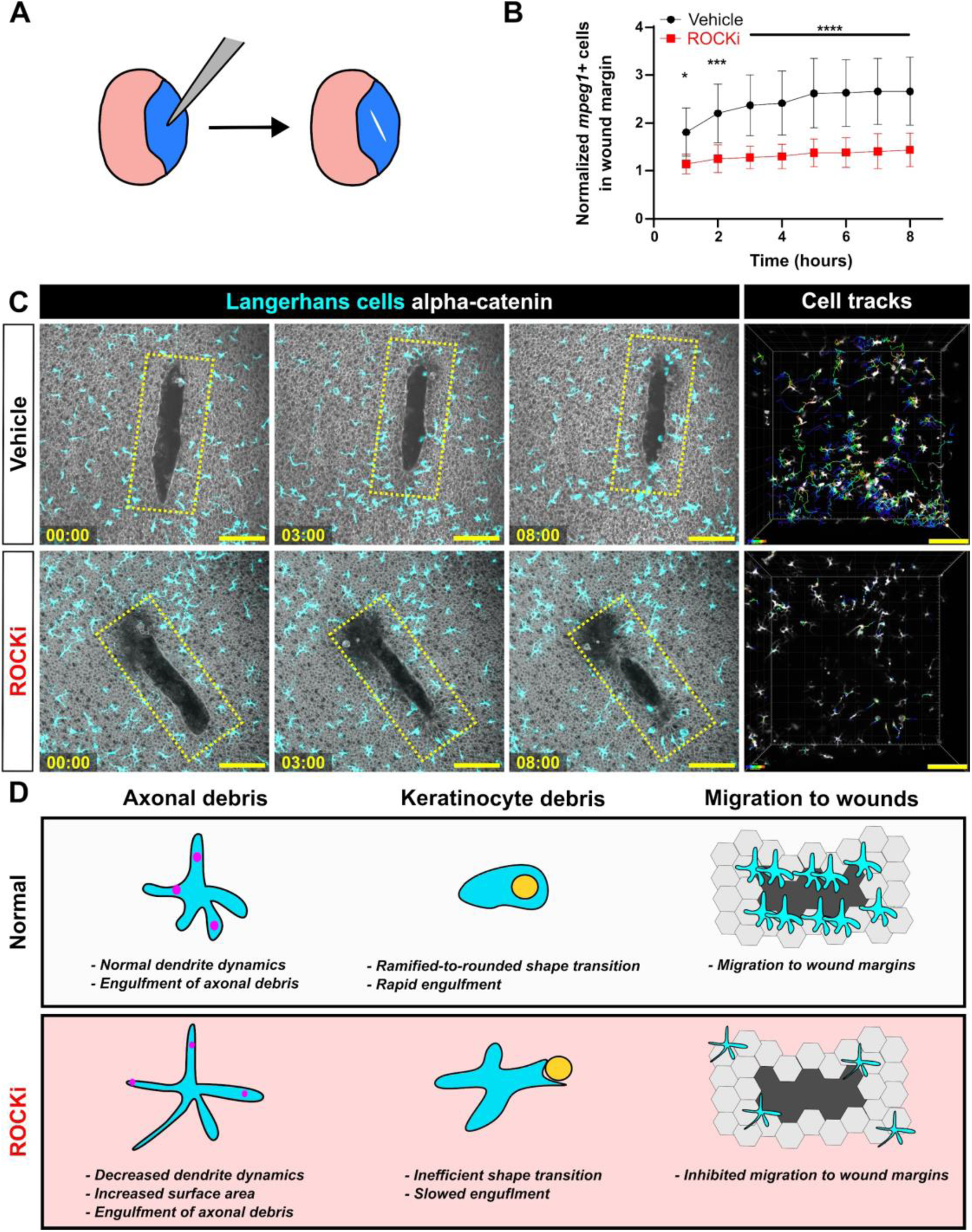
Effects of ROCK inhibition on Langerhans cell migration and graphical summary. **A.** Experimental schematic of scale scratch assay. A mechanical injury damages a swath of epidermal cells. **B.** Quantification of number of *mpeg1+* cells in wound margin, normalized to time of injury. n = 10 scales for vehicle, n = 12 scales for ROCKi. **C.** Still images of explanted scales expressing *Tg(mpeg1:mCherry)* and *Gt(ctnna1-Citrine)* depicting effects of vehicle treatment (top row) and ROCKi (bottom row). Yellow ROI denotes the wound margin used to quantify the number of *mpeg1+* cells in **(B)**. Cell tracking (rightmost panel) shows migratory tracks of vehicle- and ROCKi-treated cells color-coded according to time post-injury. **D.** Graphical summary of the effects of ROCK inhibition on Langerhans cell morphology, debris engulfment, and migration. * = p < 0.05, *** = p < 0.001, **** = p < 0.0001. Two-way ANOVA followed by Bonferroni post-tests was used to determine significance between groups at each time point in **(B)**. In **(B)**, data points represent averages, error bars represent standard deviation. Timestamps in **(C)** denote hh:mm. Scale bars in **(C)** denote 100 microns.

## DISCUSSION

Although classified as tissue-resident macrophages based on ontogeny, roles for Langerhans cells *in situ* within the epidermis remain poorly described. Here, we illustrate the dynamics and plasticity of Langerhans cells in response to multiple types of skin damage **(Figure 6D)**. During homeostasis, we show that zebrafish Langerhans cells use dynamic dendrites to surveil the tissue microenvironment, consistent with previous studies in mice.^15, 21^ Upon somatosensory axon damage and subsequent degeneration, small axonal debris appears, which Langerhans cells readily engulf with no apparent change in cell morphology. By contrast, Langerhans cells undergo a ramified-to-rounded shape transition in response to precise damage to neighboring keratinocytes. During this transition, Langerhans cell retract trailing dendrites in favor of a rounded shape more amenable to large debris engulfment. Upon acute ROCK inhibition, Langerhans cell dendrites hyper-elongate, resulting in larger areas covered but slower overall dynamics. This perturbs the ability of Langerhans cells to undergo the ramified-to-rounded shape transition and slows the engulfment process of larger debris. At the tissue-level, we show that ROCK promotes Langerhans cell migration to sites of wounds. Together, our work demonstrates that Langerhans cells are sentinels of local epidermal damage and implicates ROCK signaling as a key modulator of Langerhans cell dynamics and plasticity.

### ROCK, cellular protrusions, and complex cell shapes

Many cells elaborate specialized actin-rich protrusions, such as filopodia, microvilli, or dendrites. The length of these structures is precisely controlled by regulation of the actin cytoskeleton. For example, during filopodial retraction, many studies propose an adhesion-based feedback loop, where filopodial connections to the extracellular matrix or artificial substrates promote retrograde actin flow, causing retraction.^51–55^ Recent work concerning microvilli suggests that Myosin-IIC promotes microvilli retraction.^56^ The large dendrites possessed by Langerhans cells are distinct from both microvilli and filopodia: they are thicker than filopodia and longer than microvilli. Despite being one of the defining features of Langerhans cells, little is known about the molecular mechanisms underlying Langerhans cell dendrite morphology and dynamics. Using a Lifeact reporter, we found that F-actin dynamically remodeled during dendrite morphogenesis and debris engulfment. We showed that ROCK inhibition simultaneously slowed dendrite dynamics and increased dendrite length. Consistent with this, myosin inhibition slowed dendrite dynamics and prevented their retraction, but did not increase their lengths. Further studies are needed to decipher a precise role for ROCK substrates in the regulation of dendrite length.

Diverse cell types, including dendritic cells, microglia, astrocytes, oligodendrocytes, and neurons, share morphological similarities with Langerhans cells, possessing long cellular dendrites capable of interacting with their microenvironment. Of these, microglia are most similar functionally to the roles of Langerhans cells described herein, acting as CNS-resident macrophages capable of clearing neuronal debris and pruning synapses.^57^ Akin to our findings, an *in vitro* study found that microglial phagocytosis of apoptotic neuronal bodies required ROCK.^48^ However, in contrast to our data, an *in vivo* study found that ROCK inhibition decreased microglial surface area and dendrite number.^58^ This same study showed that ROCK inhibition also decreased microglial-neuron contacts, possibly leading to a decrease in microglia-mediated neuron elimination. It is worth noting that this study examined the brain parenchyma, in which microglia operate in a less confined 3D space compared to the densely packed epithelial environment surrounding Langerhans cells. These diverse cellular environments likely impose differential requirements for cytoskeletal effectors between tissue-resident macrophage populations. Several other cell types such as neurons and oligodendrocytes also display a ramified, protrusive morphology that aids in their biological processes. Tissue culture models showed that ROCK suppresses dendrite extension,^59^ myelin sheath formation,^60, 61^ and neurite growth^62^ in dendritic cells, oligodendrocytes, and neural stem cells, respectively. These results are consistent with our own, suggesting that ROCK regulates large cellular dendrites, thereby dictating cell morphology and function.

### Tissue-level surveillance and functions of skin-resident macrophages

Langerhans cells must achieve a balance between dendrite morphology, dynamics, and spacing to efficiently surveil the skin. A previous study using explanted skin grafts and *in vivo* imaging showed that in the absence of stimulation, Langerhans cell dendrites underwent cyclical extensions and retractions,^21^ reminiscent of our own findings. Recent *in vivo* imaging showed Langerhans cells require the small GTPase Rac1 for even spatial distribution before and after tissue tissue injury, possibly through modulating cell migration or dendrite density.^20^ The most commonly associated function with Langerhans cells is their ability to encounter and uptake antigen or pathogens and drain to lymph nodes to evoke adaptive immune responses.^7, 19, 21, 63^ The protrusive behaviors and spacing requirements reflect this: the chance that an antigen or pathogen will be encountered is increased if regular surveillance and spacing is achieved. Via our LatB treatment regimen, we found that regular dendrite behavior was necessary for engulfment of axonal debris. Our ROCKi treatment suggests a balance between dendrite motility and length in engulfment of axonal debris: an increase in dendrite length and surface area offset a decrease in dendrite dynamics.

A recent thorough analysis of macrophages in 3D matrigel found that ROCK inhibition resulted in reduced migration speeds and increased protrusiveness. Despite these alterations, ROCK-inhibited macrophages engulfed similar amounts of 3 µm latex beads as controls.^50^ Our findings regarding axonal debris engulfment after ROCK inhibition are congruent with these prior results, as we also observed an equivalent ability to engulf small debris (<∼3 µm) when compared to control cells. However, in the context of larger debris, we show that ROCK is required for efficient shape transition and engulfment by Langerhans cells. Combined with our data that ROCK promotes dendrite dynamics, we suggest that dendrite retraction is required for the rapid shape change that coincides with engulfment of larger debris. A possible corollary is that retraction of dendrites relocalizes subcellular structures, such as membrane, organelles, or cytoskeleton, to facilitate engulfment. Upon ROCK inhibition, these structures may not be redistributed in a timely fashion, resulting in slowed engulfment. Future experiments testing redistribution of these components will be required to fully ascertain the role for ROCK in phagocytosis of large, but not small, debris.

What are the functions for macrophages in response to tissue-level injuries? Recent elegant work described a pro-angiogenic requirement for Langerhans cells after large wounds in mice.^4^ Beyond skin-resident macrophages, a plethora of work has examined the roles of *D. melanogaster* hemocytes (macrophage-like cells) in embryonic epithelial wound repair. Most notably, and consistent with our results, inhibiting the small GTPase *rho*, which functions immediately upstream of ROCK, prevents hemocyte migration to wound sites.^64^ In vertebrate systems, depletion of entire macrophage populations can lead to wildly different results, depending on the exact model used, ablation method, and timing.^65–69^ In our study, we found that inhibiting ROCK led to slower debris engulfment dynamics by Langerhans cells. Further, we showed that inhibiting ROCK prevented Langerhans cell migration to large tissue-scale wounds. Whether or not slowed engulfment of debris and migration by Langerhans cells impedes efficient wound healing over a period of days remains to be determined.

Overall, our work highlights the power of the zebrafish system for analysis of the rapid and plastic responses of Langerhans cells to acute epidermal perturbations. This study reveals a critical role for ROCK signaling in the ability of skin-resident macrophages to dynamically surveil the epidermis and respond to debris and injuries of different magnitudes.

## Acknowledgements

We thank the LSB Aquatics staff for animal care, Dan Fong and Wai Pang Chan for imaging support. We thank Anna Huttenlocher’s lab for providing the *mpeg1:Lifeact-mRuby* plasmid, Julie Theriot’s lab for helpful discussion and providing reagents, and Alvaro Sagasti and Francisco Barros-Becker for helpful discussion. The authors are grateful to all members of the Rasmussen lab for critical support, critiques and discussion.

## Funding

This investigation was supported by a Washington Research Foundation Postdoctoral Fellowship to E.P., an award from the Fred Hutchinson Cancer Research Center/University of Washington Cancer Consortium (P30 CA015704) and funds from the University of Washington to J.P.R. J.P.R. is a Washington Research Foundation Distinguished Investigator.

## Author contributions

Conceptualization: E.P.,J.P.R.; Methodology: E.P., E.J.A.Q., J.P.R.; Formal analysis: E.P., E.J.A.Q., C.E.A.G.; Investigation: E.P.; Resources: J.P.R.; Writing-original draft: E.P.; Writing-review & editing: E.P., J.P.R.; Visualization: E.P.,J.P.R.; Supervision: E.P.,J.P.R.; Project administration: E.P.,J.P.R.; Funding acquisition: E.P.,J.P.R.

## Declaration of interests

The authors declare no competing interests.

## MATERIALS AND METHODS

### Key resources table

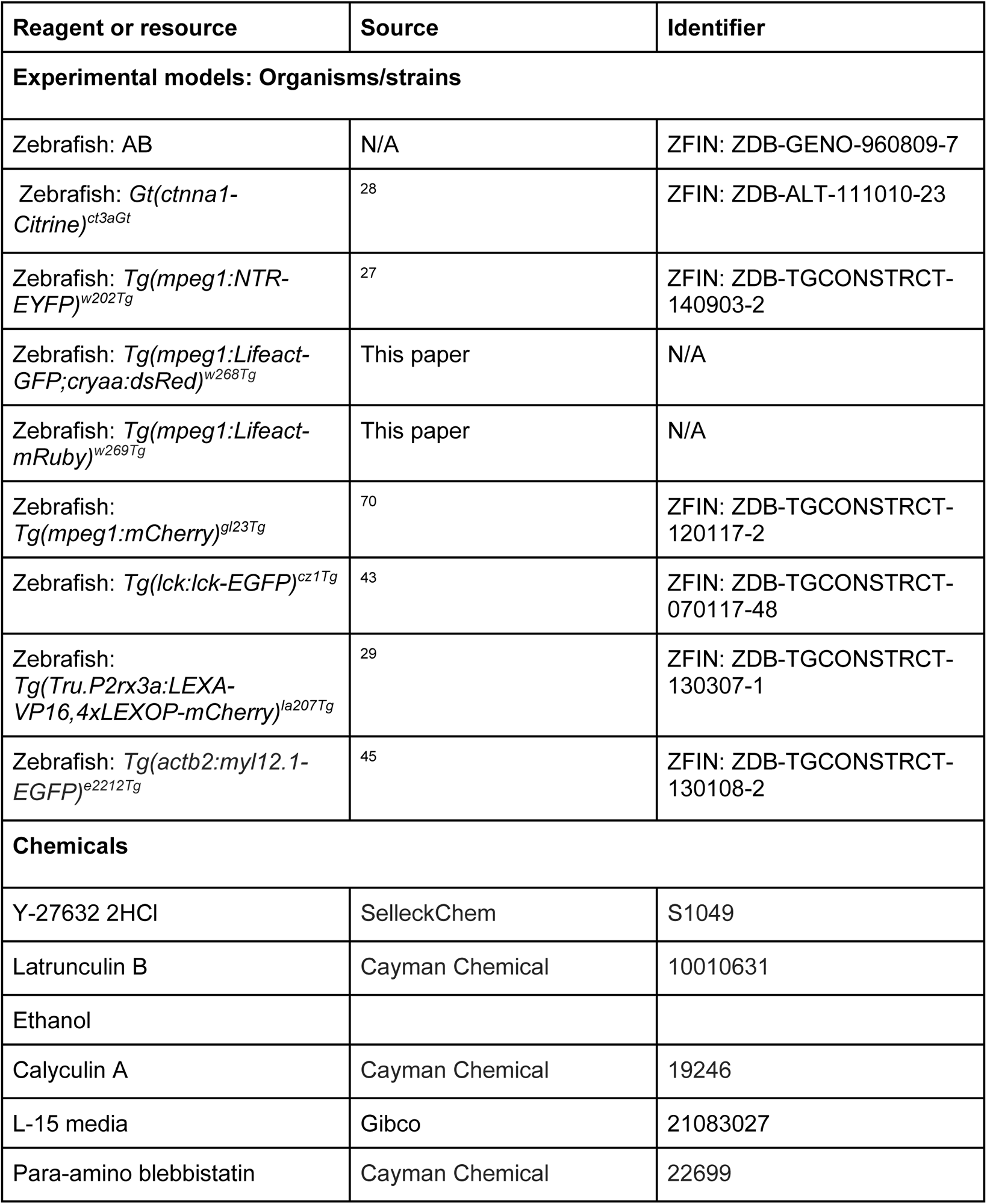
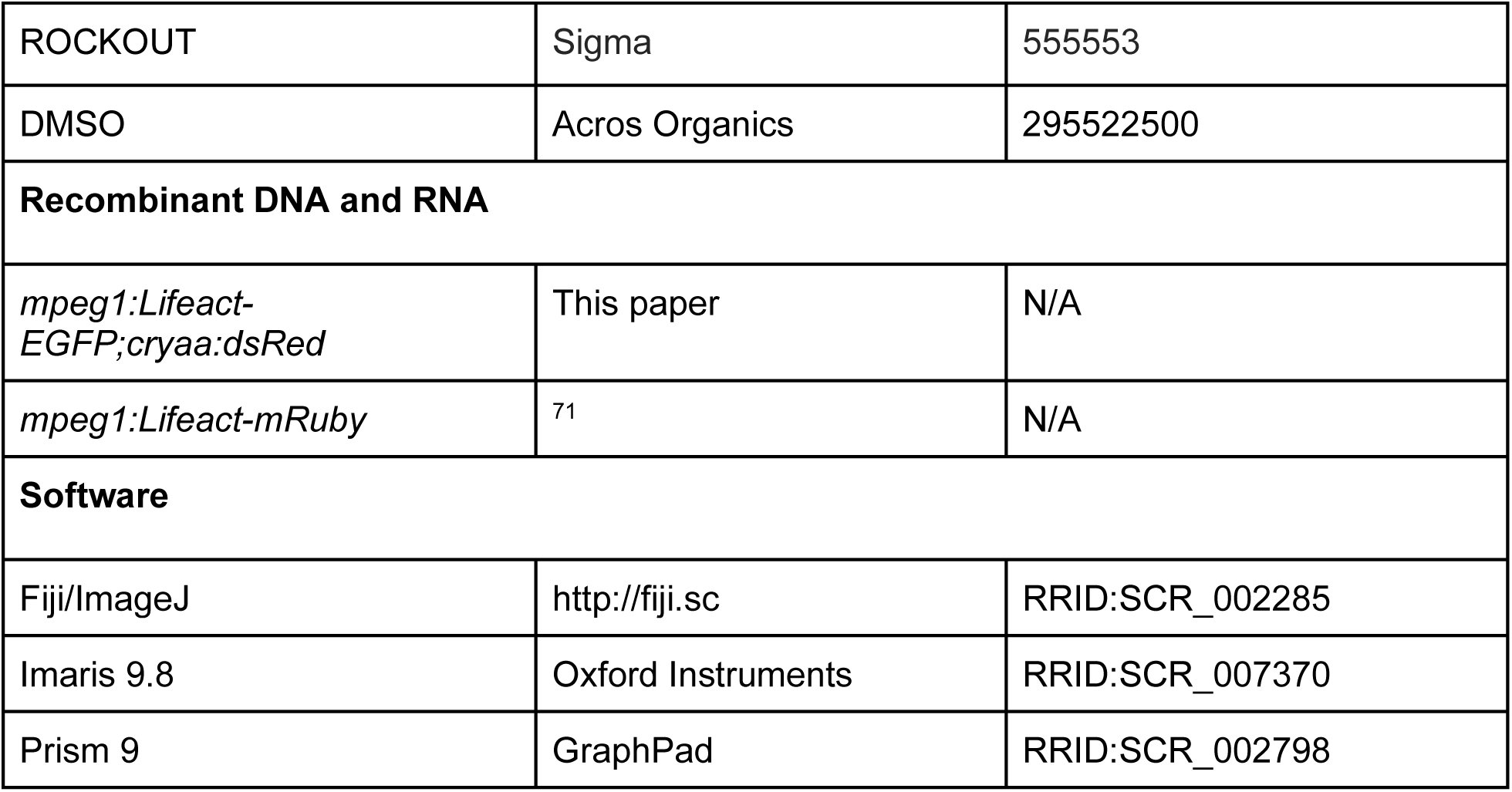

### Zebrafish husbandry

Zebrafish were housed at 26-27°C on a 14/10 h light cycle. The strains used are listed in the Key Resources Table. Animals aged 6-18 months of either sex were used in this study. All zebrafish experiments were approved by the Institutional Animal Care and Use Committee at the University of Washington (Protocol #4439-01).

### Generation of transgenic zebrafish

To generate *Tg(mpeg1:Lifeact-EGFP;cryaa:DsRed)^w^*^268*Tg*^, Gibson assembly was used to create a vector containing *tol2* arms and the *mpeg1.1* promoter driving Lifeact-EGFP. The *alpha A crystallin* (*cryaa*) promoter driving DsRed was used as a transgenesis marker. Wildtype zebrafish embryos were injected with plasmid DNA and *tol2* mRNA at the 1-cell stage, and embryos with the red lens marker were raised to adulthood. Adults were initially screened for GFP+ cells in the skin, and GFP+ adults were then outcrossed to wild-type partners. F1 fish with red lens markers were raised to adulthood, where GFP expression was assessed.

To generate *Tg(mpeg1:Lifeact-mRuby)^w^*^269*Tg*^, the previously published *mpeg1:Lifeact-mRuby* plasmid from Barros-Becker et al.^71^ and *tol2* mRNA were injected to wild-type embryos at the 1-cell stage. Adults were screened for mRuby+ cells in the skin. mRuby+ adults were then outcrossed to wild-type partners. F1 fish were raised to adulthood, where mRuby expression was assessed.

### Scale removal and scale injury assay

For scale removal, adult fish were anesthetized in system water containing 200 µg/ml buffered tricaine, and forceps were used to remove individual scales. Following scale removal, animals were recovered in system water.

For the scale injury assay in Figure 6, scales were explanted and treated for 40 minutes with DMSO or Y-27632. After 40 minutes, scales were placed under a dissecting microscope. One pair of forceps was used to assist in pinning the scale down by contacting a region devoid of epidermis. A second pair introduced the scratch in the middle of the epidermis (as depicted in Figure 6A). Scratches were only used for data collection if they did not extend to the edge of the scale and were oval in shape (as shown by representative images in Figure 6B).

### Microscopy and live imaging

An upright Nikon Ni-E A1R MP+ confocal microscope was used for all experiments. A 25× water dipping objective (1.1 NA) was routinely used. Unless otherwise stated, scales were removed and placed onto dry 6 mm plastic dishes, epidermis side up, and allowed to adhere for 1 min before adding L-15 medium pre-warmed to room temperature. For experiments involving axon degeneration, scales were incubated at 26°C for 90-120 min followed by imaging, which was performed at room temperature (23°C).

### Chemical treatments

For Latrunculin B (Cayman Chemical, 10010631), Y-27632 2HCl (SelleckChem S1049), ROCKOUT, (Sigma, 555553), para-amino blebbistatin (Cayman Chemical, 22699) and calyculin A (Cayman Chemical, 19246) treatments, scales were removed and immediately placed in L-15 media. Imaging commenced at least 15 minutes before careful addition of chemicals while on the microscope stage. Final concentrations used in this study: Latrunculin B, 10 µM; Y-27632, 50 µM; ROCKOUT, 100 µM; para-amino blebbistatin, 100 µM; and calyculin A, 250 nm. Appropriate vehicle controls, either DMSO or ethanol, were used at equivalent %v/v.

For washout experiments, Latrunculin B or Y-27632 was added to 5 ml of L-15 and added to a dish of explanted scales. After 5 (LatB) or 20 (Y-27632) minutes, media was exchanged 4 times using 5 ml for each wash. The dish was immediately placed onto the microscope stage and imaging commenced.

### Laser-induced cell damage

For laser-induced cell damage, scales were mounted into the imaging chamber as described above. Target cells at least 1 cell distance away from a Langerhans cell (∼5-15 microns) and within the same z-plane were located and ablated using a UGA-42 Caliburn pulsed 532 nm laser (Rapp OptoElectronic). The laser was focused through a 25× objective at 4× zoom. Ablation was produced in the focal plane using 15-20% power at a single point within a nucleus, firing 3 times for 3 seconds each using a custom NIS-Elements macro.

### Image analysis

The Imaris Filaments package was used to skeletonize cells and track individual dendrite dynamics (Figures 1D and 4D, E). Cumulative sum distances traveled by dendrites were calculated by summing a random 10 minute tracked segment. The Imaris “Surfaces” function was used to calculate the volume of debris engulfed as previously performed.^13^ Individual cells were traced within ImageJ to track and calculate convex hull and circularity. dendrite number (greater than 5 microns) and lifetime were manually counted. To calculate dendrite and retraction speeds, individual dendrites were skeletonized; the average speed of extension or retraction across multiple frames was calculated and graphed. To calculate “Time to First Contact” in Figure 5F, the image from the transmitted detector channel was used as a reference to visualize larger debris. The fluorescence image was overlaid onto the transmitted detector image and cells were tracked as they encountered debris post-ablation. The difference between time of ablation and time of first contact represents the data in Figure 5F. To calculate “Time from contact to closure” the difference in time from contact to full closure of the phagocytic cup was calculated and represents the data in Figure 5G. These same time points were used as reference time points for calculating circularity and surface area at time of engulfment in Figure 5D, E. To calculate the cell number in wound margin in Figure 6C, an ROI of 150 microns wide and (length of wound x 1.2) microns long was drawn around the wound. The number of cells was counted for each timepoint and normalized to time 0. Cells were only considered if >50% of their cell body was within the ROI. Cell tracking was performed using the “Cells” function in Imaris.

To generate *Tg(actb2:myl12.1-EGFP)* images in Supplemental Figure 4, Imaris was used to generate masks of the red channel (*Tg(mpeg1:Lifeact-mRuby)*). Then, GFP+ signal within these masks was used to generate the dendrite-specific NMII signal seen in Supplemental Figure 4A.

### Statistical analysis

GraphPad Prism was used to generate graphs and perform statistical analyses. At least three individual biological experiments were performed unless otherwise noted. Tests used and number scales or cells/ROIs are described in each figure legend.

## SUPPLEMENTAL FIGURES

**Supplemental Figure 1.**
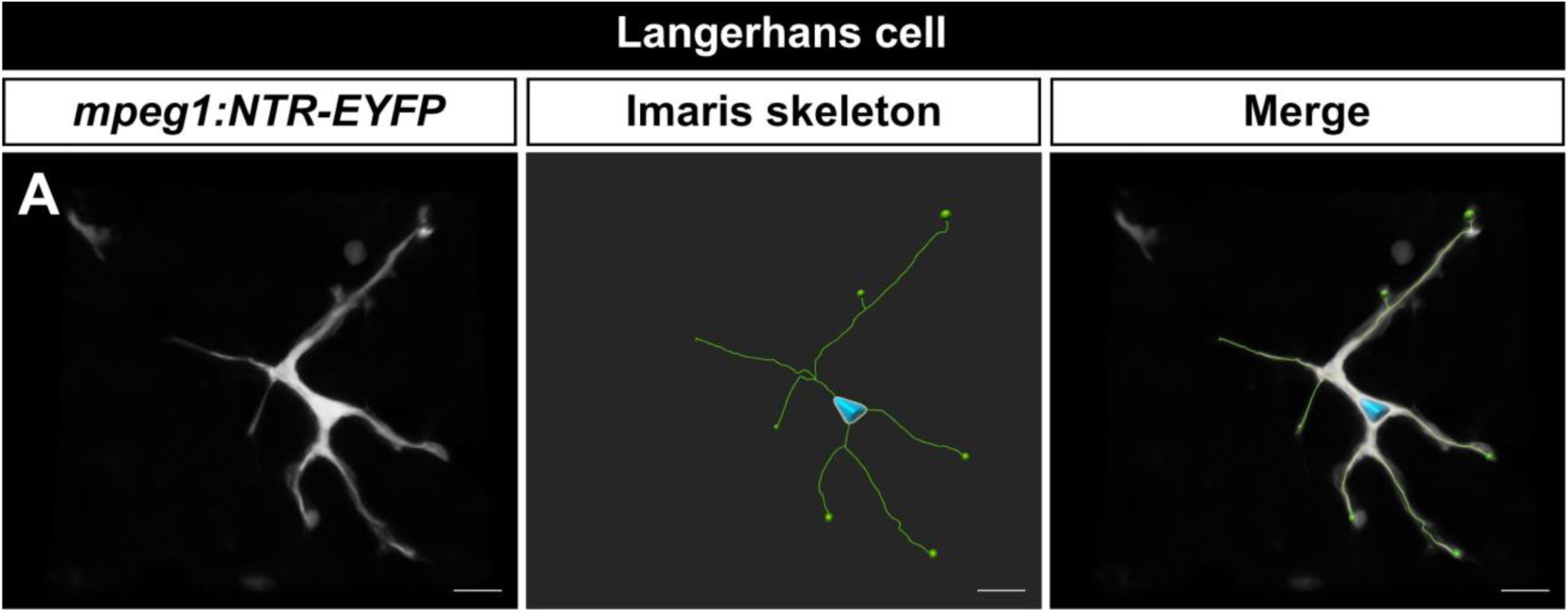
Skeletonization of a Langerhans cell. **A.** Representative image of *Tg(mpeg1:NTR-EYFP)* skeletonized using Imaris Filaments module. See also Supplemental Video 1. Scale bar in **(A)** denotes 10 microns.

**Supplemental Figure 2.**
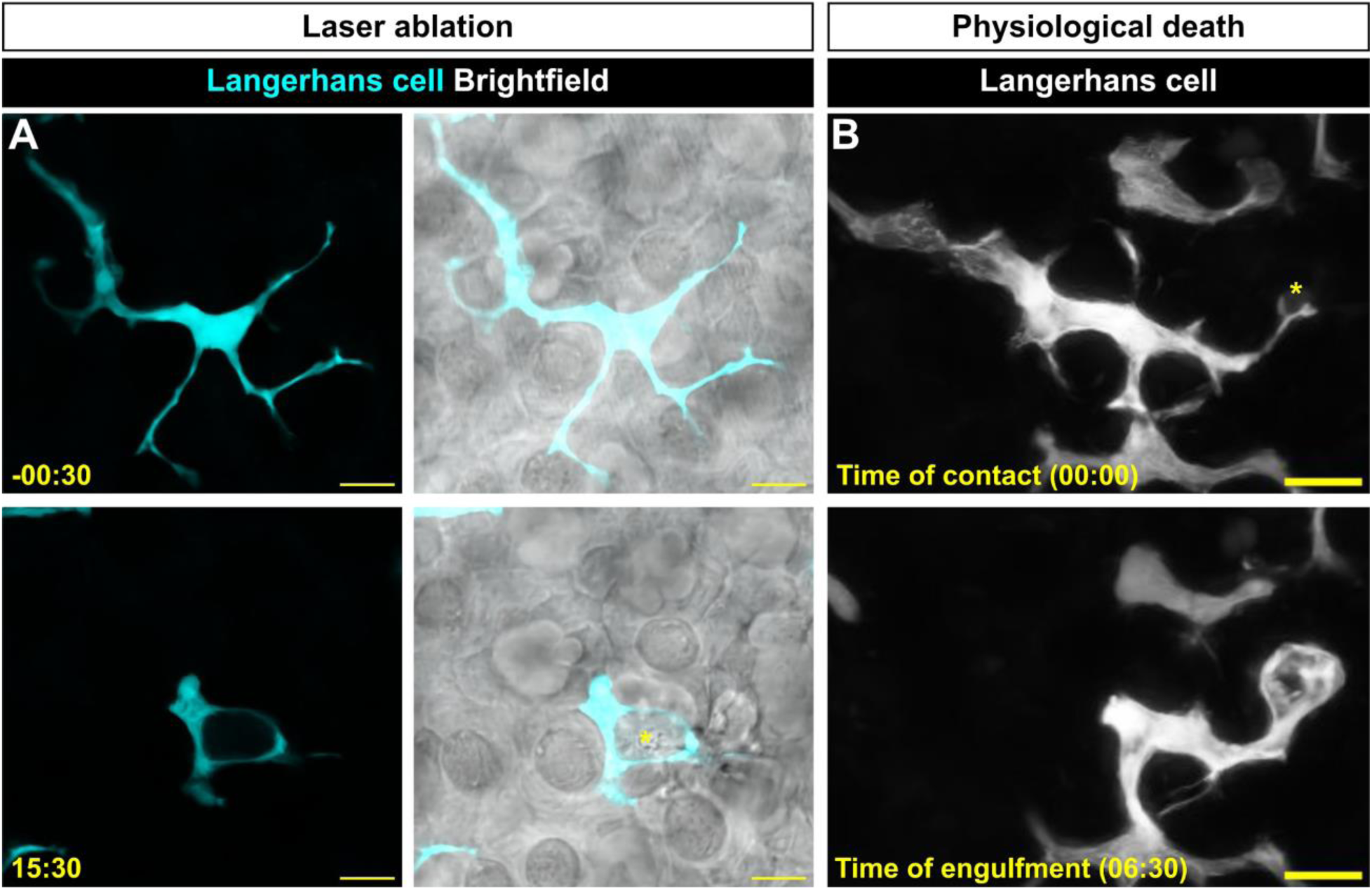
Langerhans cells undergo shape changes during engulfment of large cellular debris. **A.** Stills of *Tg(mpeg1:NTR-EYFP)*-positive Langerhans cell showing fluorescent only (left) and fluorescent+brightfield composite (right) images before and after laser ablation. Asterisk depicts location of laser ablation. **B.** Stills of *Tg(mpeg1:NTR-EYFP)* showing “natural” engulfment of large debris in the absence of laser ablation. Asterisk depicts future site of phagocytosis. Timestamps denote mm:ss. Scale bars denote 10 microns.

**Supplemental Figure 3.**
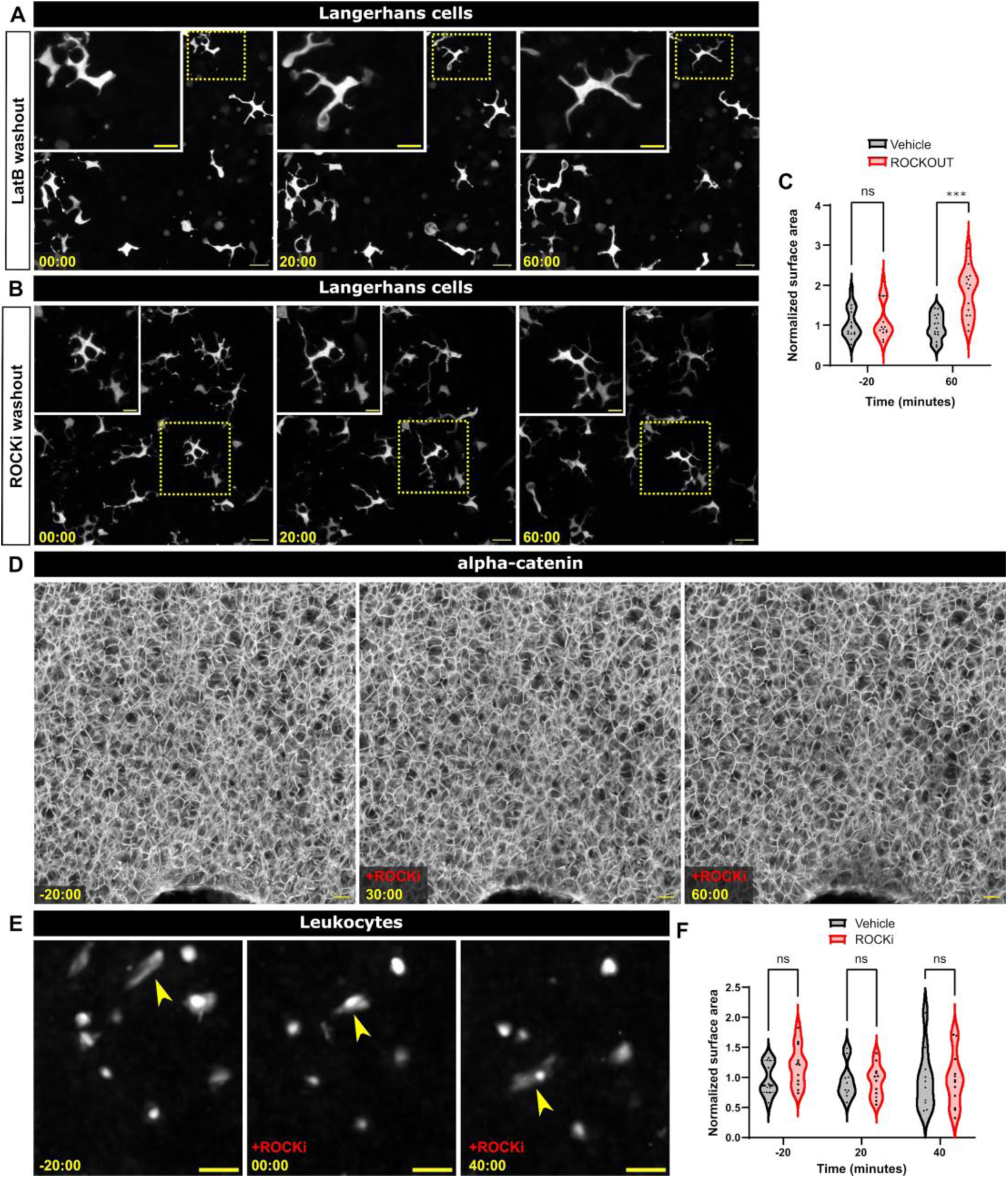
Effects of LatB and ROCK inhibition of Langerhans cells. **A.** Stills of *Tg(mpeg1:NTR-EYFP)*-positive Langerhans cells depicting normal cell motility after LatB washout. **B.** Stills of *Tg(mpeg1:NTR-EYFP)*-positive Langerhans cells depicting normal cell motility after Y-27632 washout. Dotted boxes in **(A,B)** denote regions magnified in insets. **C.** Quantification of surface area covered after ROCKOUT treatment, an alternative inhibitor of ROCK. Data shown are compiled from two individual experiments, n = 15 cells from N = 6 scales for vehicle treatment and n = 14 from N = 5 scales for ROCKOUT treatment. **D.** Stills of *Gt(ctnna1-Citrine)*-positive cells depicting normal tissue morphology after ROCK inhibition. **E.** Stills of *Tg(lck:lck-GFP)*-positive leukocytes showing no changes in cell morphology after ROCK inhibition. **F.** Violin plots of surface area covered by *Tg(lck:lck-GFP)*-positive cells after ROCKi. Data shown are compiled from two individual experiments, n = 10 cells from N = 5 scales for vehicle treatment and n = 12 from N = 4 scales for ROCki treatment.*** = p < 0.0001. Mann-Whitney U tests were used to determine significance between groups at each time point in **(C)**. Two-way ANOVA followed by Bonferroni correction was used in **(F)**. Timestamps denote mm:ss. Scale bars in **(A, B, D)** denote 20 microns, scale bars in **(A, inset; B, inset; E)** denote 10 microns.

**Supplemental Figure 4.**
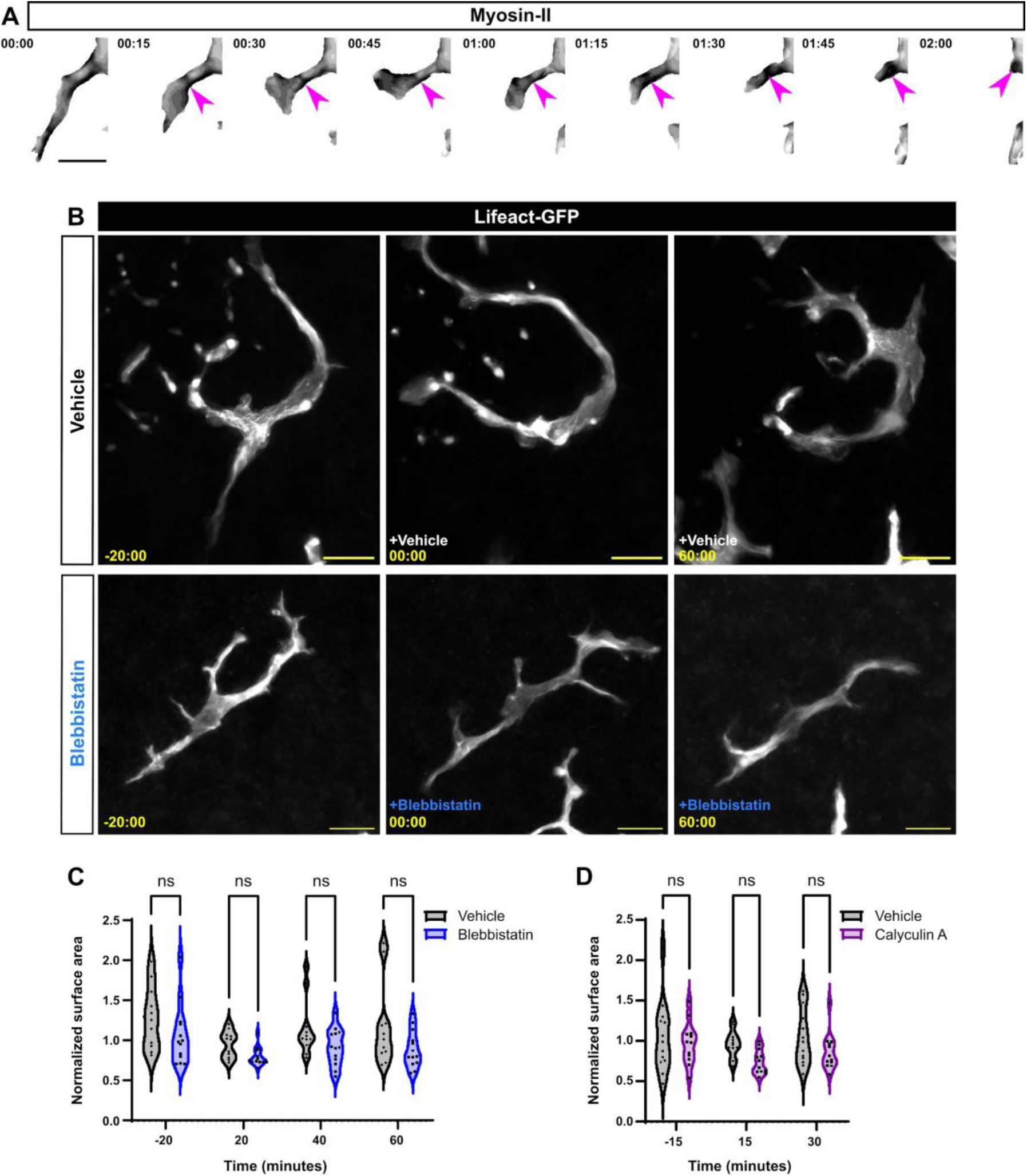
Myosin perturbation and its effects on Langerhans cell shape. **A.** Stills of macrophage-specific myosin (inverted grayscale) during dendrite retraction (see Materials and Methods for details). Magenta arrowheads indicate higher levels of myosin during dendrite retraction. **B.** Stills of *Tg(mpeg1:Lifeact-GFP)*-positive cells treated with vehicle **(B, top)** or para-amino blebbistatin **(B, bottom)**. **C.** Violin plots of Langerhans cell surface area, normalized to time of treatment with vehicle or para-amino blebbistatin. Data are representative of two individual experiments, n = 11 from N = 4 scales cells for vehicle treatment and n = 13 cells from N =3 scales for blebbistatin treatment. **D.** Violin plots of Langerhans cell surface area, normalized to time of treatment with vehicle or Calyculin A. Data are representative of two individual experiments, n = 13 cells from N = 6 scales for vehicle treatment and n = 12 cells from N = 5 scales for Calyculin A treatment. Two-way ANOVA followed by Bonferroni post-tests revealed no significant differences. Timestamps in B denote mm:ss. Scale bars in **(A, B)** denote 10 microns.

## SUPPLEMENTAL VIDEO LEGENDS

**Supplemental Video 1.** Time-lapse microscopy of Langerhans cell (cyan) extending and retracting protrusions among epidermal cell membranes (white). Video was bleach-corrected. Scale bar denotes 10 microns.

**Supplemental Video 2.** Time-lapse microscopy of Langerhans cell (cyan) engulfing *Tg(p2rx3a:mCherry)*+ debris (magenta). Scale bar denotes 10 microns.

**Supplemental Video 3.** Time-lapse microscopy of Langerhans cell (white) engulfing cellular debris generated after laser-induced damage of keratinocytes. Yellow asterisk denotes site of ablation. Scale bar denotes 10 microns.

**Supplemental Video 4.** Time-lapse microscopy of Langerhans cell labeled with Lifeact-EGFP. Scale bar denotes 10 microns.

**Supplemental Video 5.** Time-lapse microscopy of Langerhans cell labeled with Lifeact-EGFP (false-colored) engulfing *Tg(p2rx3a:mCherry)*+ debris (white). Scale bar denotes 10 microns.

**Supplemental Video 6.** Time-lapse microscopy of Langerhans cell labeled with Lifeact-EGFP (false-colored) engulfing cellular debris generated after laser-induced damage of keratinocytes. Yellow asterisk indicates site of ablation. Red arrowhead indicates Lifeact-EGFP accumulation during engulfment. Scale bar denotes 10 microns.

**Supplemental Video 7.** Time-lapse microscopy of Langerhans cells (white) treated with vehicle or ROCK inhibitor. Scale bar denotes 20 microns.

**Supplemental Video 8.** Time-lapse microscopy of Langerhans cell (white) engulfing cellular debris generated after laser-induced damage of keratinocytes. Cells are treated with vehicle or ROCK inhibitor. Scale bar denotes 10 microns.

**Supplemental Video 9.** Time-lapse microscopy of Langerhans cells (cyan) reacting to epidermal wounds (epidermal cells labeled in white). Cells are treated with vehicle or ROCK inhibitor. Scale bar denotes 100 microns.

## Notes

### Competing Interest Statement

The authors have declared no competing interest.

